# Different efficacies of neutralizing antibodies and antiviral drugs on SARS-CoV-2 Omicron subvariants, BA.1 and BA.2

**DOI:** 10.1101/2022.02.27.482147

**Authors:** Hirofumi Ohashi, Takayuki Hishiki, Daisuke Akazawa, Kwang Su Kim, Joohyeon Woo, Kaho Shionoya, Kana Tsuchimoto, Shoya Iwanami, Saya Moriyama, Hitomi Kinoshita, Souichi Yamada, Yudai Kuroda, Tsukasa Yamamoto, Noriko Kishida, Shinji Watanabe, Hideki Hasegawa, Hideki Ebihara, Tadaki Suzuki, Ken Maeda, Shuetsu Fukushi, Yoshimasa Takahashi, Shingo Iwami, Koichi Watashi

## Abstract

The severe acute respiratory syndrome coronavirus 2 (SARS-CoV-2) Omicron subvariant BA.2 has spread in many countries, replacing the earlier Omicron subvariant BA.1 and other variants. Here, using a cell culture infection assay, we quantified the intrinsic sensitivity of BA.2 and BA.1 compared with other variants of concern, Alpha, Gamma, and Delta, to five approved-neutralizing antibodies and antiviral drugs. Our assay revealed the diverse sensitivities of these variants to antibodies, including the loss of response of both BA.1 and BA.2 to casirivimab and of BA.1 to imdevimab. In contrast, EIDD-1931 and nirmatrelvir showed a more conserved activities to these variants. The viral response profile combined with mathematical analysis estimated differences in antiviral effects among variants in the clinical concentrations. These analyses provide essential evidence that gives insight into variant emergence’s impact on choosing optimal drug treatment.

## Introduction

The severe acute respiratory syndrome coronavirus 2 (SARS-CoV-2) Omicron variant (lineage B.1.1.529) has rapidly spread worldwide and become the most prevalent SARS-CoV-2 in many countries (Elliott et al., 2022; Viana et al., 2022). Of the identified Omicron subvariants, the subvariant BA.1 was dominantly prevalent in the early days after Omicron emerged from November 2021. However, the replacement of BA.1 with another subvariant, BA.2, has grown in the countries, including Denmark, UK, and South Africa, alerting a higher transmission of this new subvariant worldwide that can prolong the current wave of COVID-19 (USKH, 2022). The BA.1 and BA.2 have more than 30 shared amino acid substitutions from the Wuhan strain, especially with approximately 20 shared mutations in the Spike protein. They also have some unique mutations (Figure 1). For example, the S1 69-70 deletion as a hallmark of BA.1, associated with S-gene target failure in PCR tests, is unconserved in BA.2 (WHO, 2021; Majumdar and Sarkar, 2021). BA.2 also has four unique substitutions (S371F, T376A, D405N, and R408S) compared with BA.1, with lacking three mutations (S371L, G446S, and G496S) in the receptor-binding domain of the S1, which is involved in vaccine and antibody responses (Majumdar and Sarkar, 2021). Such unique mutation patterns in BA.1 and BA.2 possibly affect their sensitivities to approved drugs/antibodies against COVID-19. Therefore, we quantified such drug/antibody responses of BA1 and BA.2 compared to other variants of concern (Alpha, Gamma, Delta) and a Wuhan strain in cell culture infection assays. Furthermore, most reports have so far evaluated only 50% (or 90%) inhibitory concentrations to quantify the drug activity. Yet, these concentrations are pharmacologically not the sole factor that determines antiviral efficacy. Thus, we also estimated the slopes of dose-response sigmoid curves to quantitatively discuss their drug effects at clinical drug concentrations (Koizumi et al., 2017; Shen et al., 2008).

**Figure 1.**
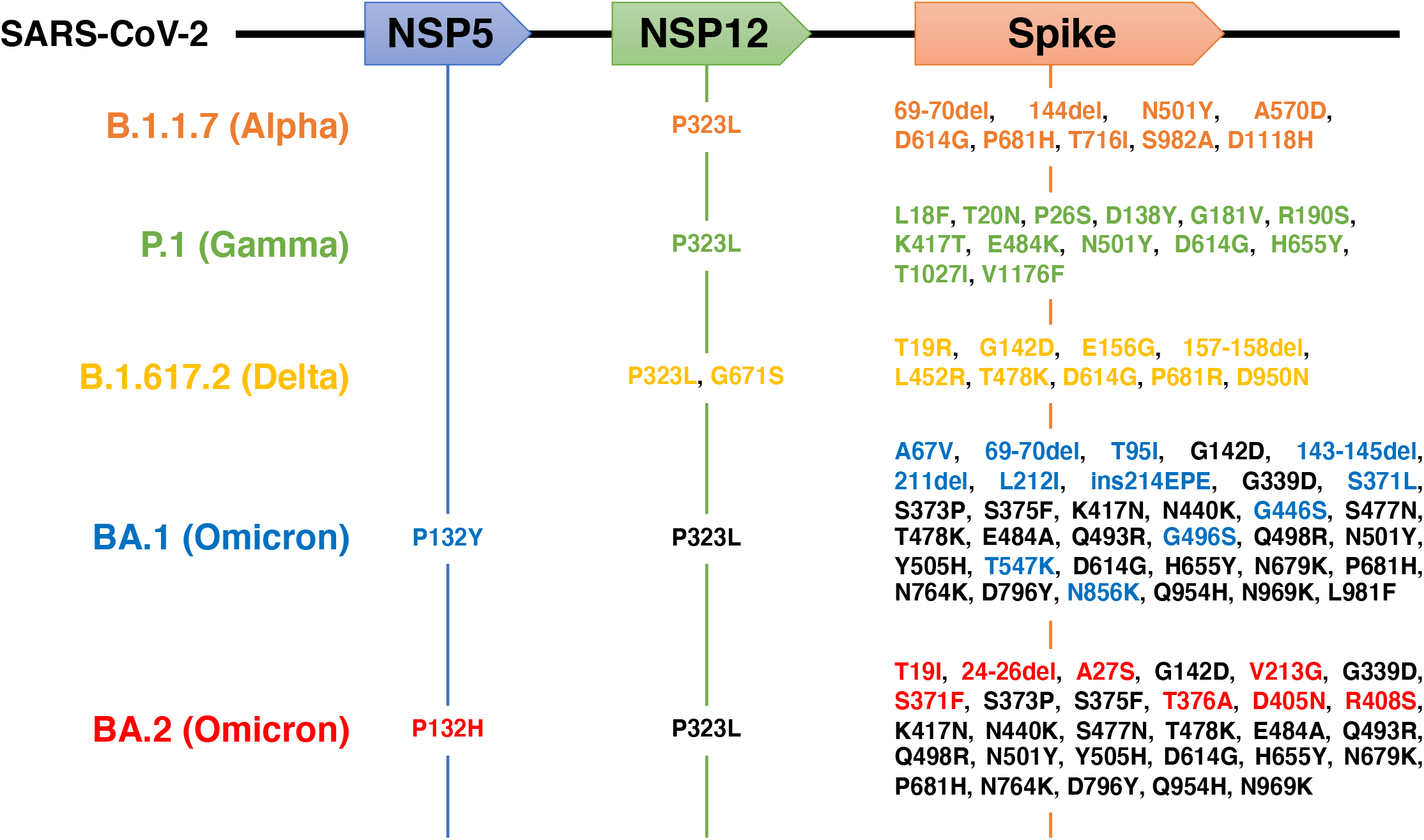
Schematic representations for amino acid substitutions within the B.1.1.7, P.1, B.1.617.2, BA.1, and BA.2 lineage in NSP5, NSP12, and Spike proteins. Upper boxes show coding regions for NSP5 (the target of nirmatrelvir), NSP12 (the target of EIDD-1931), and Spike (the target of imdevimab, casirivimab, and S309) in the SARS-CoV-2 genome RNA. Mutated amino acids from the Wuhan strain in B.1.1.7 (Alpha, orange), P.1 (Gamma, green), B.1.617.2 (Delta, yellow), BA.1 (Omicron), and BA.2 (Omicron) are shown. Shared BA.1 and BA.2 mutations are indicated in black, and those unique to BA.1 and BA.2 are shown in blue and red, respectively.

## Results and Discussion

### Dose response curves of approved-antibodies/drugs on SARS-CoV-2 variants

We evaluated the intrinsic sensitivity of SARS-CoV-2 variants (Wuhan, Alpha, Gamma, Delta, Omicron-BA.1, and Omicron-BA.2) to the approved antibodies/drugs [casirivimab, imdevimab, S309 (the prototype antibody of sotrovimab), EIDD-1931 (the active form of molnupiravir), and nirmatrelvir]. Each SARS-CoV-2 strain was inoculated and cultivated in VeroE6/TMPRSS2 cells upon treatment with varying concentrations of antibodies/drugs (up to 4-10 μM or μg/mL) to measure viral RNA in the culture supernatant, as well as cell viability at 24 h postinoculation (Matsuyama et al., 2020). Figure 2 shows the dose-response curve of each variant against tested antibodies/drugs (Figure 2). No cytotoxicity induced by antibody/drug was observed in all tested concentrations (Figure S1). Overall, inhibition potency of the three tested antibodies, casirivimab, imdevimab, and S309 to Omicron subvariants BA.1 and BA.2, were severely impaired, in contrast to their outstanding activities against the Wuhan strain and Alpha, Gamma, and Delta variants (Figure 2A-C). Casirivimab did not show any antiviral activity to BA.1 and BA.2 up to 10 μg/ml (Figure 2A). Also, imdevimab lost its activity to BA.1, but retained a minor antiviral activity to reduce BA.2 infections (Figure 2B). S309’s antiviral activity to BA.1 was more modest than that of other variants, and that to BA.2 was even weaker (Figure 2C). These tendencies of the IC50 shifts between BA.1 and other variants (Table. 1) are overall consistent with the previous reports (Cameroni et al., 2021; Cao et al., 2021; Liu et al., 2021; Planas et al., 2021). Additionally, our dose-response curves clearly show the impaired potency of all three antibodies against BA.2. As a possible mechanistic explanation, a class 2 antibody, casirivimab, completely lost its antiviral activity to both BA.1 and BA.2, probably because of the mutations at K417N, S477N, T478K, E484A, Q493R, Q498R, and N501Y (Figure 1, black), contained in the reported epitope footprints of casirivimab (VanBlargan et al., 2022). The class 3 imdevimab showed the reduced activity to BA.2 compared with the Wuhan and other variants, which can be explained by the N440K, Q498R, and N501Y mutations (Figure 1, black) within the imdevimab epitope amino acids, and were inactive to BA.1, carrying a further mutation at G446S (Figure 1, blue) within the imdevimab epitope. Another class 3 S309 also showed a reduced antiviral activity against BA.1 and BA.2, containing G339D and N440K substitutions from the Wuhan strain (Figure 1, black) within the S309 epitope footprint. In contrast to antibodies, a polymerase inhibitor, EIDD-1931, and a main protease inhibitor, nirmatrelvir, dose-dependently reduced the viral RNA of all variants and showed no resistance (Figure 2D, E), consistent with the previous reports showing no remarkable IC50 differences among BA.1 and other variants (Li et al., 2022; Takashita et al., 2022; Vangeel et al., 2022).

**Figure 2.**
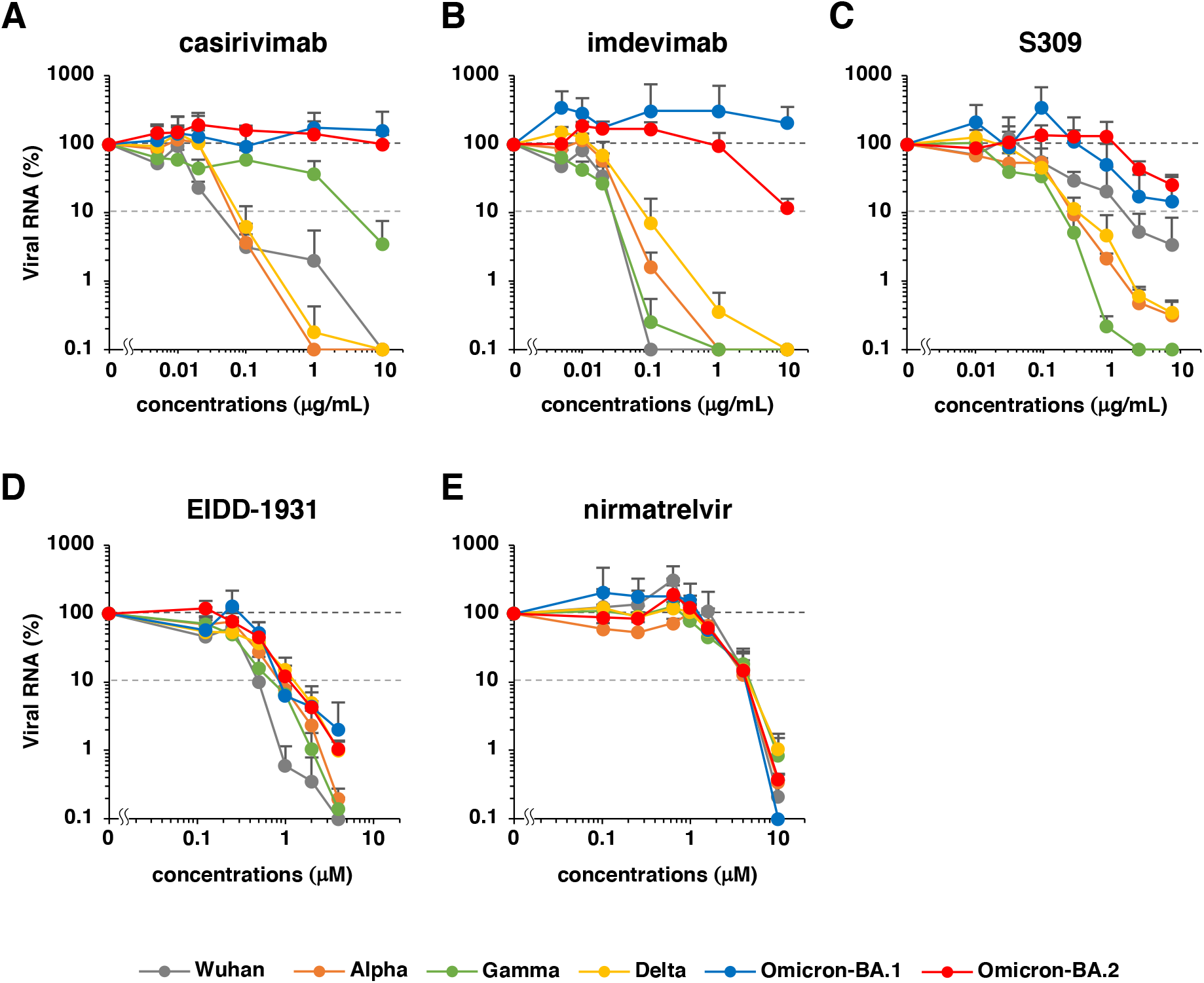
Dose-response curves for each SARS-CoV-2 variant propagation upon antibody or drug treatment. Relative SARS-CoV-2 RNAs were plotted in log-scale against the concentrations of approved antibodies/drugs [A: casirivimab, B: imdevimab, C: S309 (a parent antibody of sotrovimab), D: EIDD-1931 (the active form of molnupiravir), and E: nirmatrelvir]. Gray: Wuhan strain, orange: Alpha, green: Gamma, yellow: Delta, blue: Omicron-BA.1, and red: Omicron-BA.2. Data are presented as mean ± SD across the three replicate experiments. Relative values are shown as percentages of viral RNA in culture supernatants to the control wells incubated without antibodies/drugs. Values less than 0.1% are shown as 0.1% in these graphs.

**Table 1.** Estimated parameters for the antiviral effect of drugs

| | $IC_{50}$ | $IC_{90}$ | $m$ | $IIP (C_{max})$ |
| --- | --- | --- | --- | --- |
| **casirivimab** | ($\mu\text{g/ml}$) | ($\mu\text{g/ml}$) | | |
| WK-521 | 0.0139 | 0.0713 | 1.3444 | 5.5662 |
| QK002 (Alpha) | 0.0136 | 0.0505 | 1.6747 | 6.9496 |
| TY7-501 (Gamma) | 0.0140 | 0.1666 | 0.8872 | 3.6706 |
| TY11-927 (Delta) | 0.0217 | 0.0749 | 1.7743 | 7.0029 |
| TY38-873 (Omicron. BA.1) | > 10 | - | - | - |
| TY40-385 (Omicron. BA.2) | > 10 | - | - | - |
| **imdevimab** | ($\mu\text{g/ml}$) | ($\mu\text{g/ml}$) | | |
| WK-521 | 0.0125 | 0.0963 | 1.0763 | 4.5202 |
| QK002 (Alpha) | 0.0227 | 0.0813 | 1.7221 | 6.7862 |
| TY7-501 (Gamma) | 0.0082 | 0.0416 | 1.3529 | 5.9296 |
| TY11-927 (Delta) | 0.0290 | 0.0834 | 2.0809 | 7.9787 |
| TY38-873 (Omicron. BA.1) | > 10 | - | - | - |
| TY40-385 (Omicron. BA.2) | 1.2525 | 11.6931 | 0.9836 | 2.1658 |
| **S309** | ($\mu\text{g/ml}$) | ($\mu\text{g/ml}$) | | |
| WK-521 | 0.1587 | 0.7583 | 1.4048 | 4.0316 |
| QK002 (Alpha) | 0.0552 | 0.5449 | 0.9596 | 3.1943 |
| TY7-501 (Gamma) | 0.0384 | 0.1498 | 1.6143 | 5.6276 |
| TY11-927 (Delta) | 0.0870 | 0.2407 | 2.1589 | 6.7593 |
| TY38-873 (Omicron. BA.1) | 0.9579 | 2.6822 | 2.1348 | 4.4598 |
| TY40-385 (Omicron. BA.2) | 1.3579 | 62.2520 | 0.5744 | 1.1452 |
| **EIDD-1931** | ($\mu\text{M}$) | ($\mu\text{M}$) | | |
| WK-521 | 0.2270 | 0.7385 | 1.8626 | 2.9789 |
| QK002 (Alpha) | 0.3435 | 1.0099 | 2.0375 | 2.8922 |
| TY7-501 (Gamma) | 0.2432 | 0.6531 | 2.2241 | 3.4901 |
| TY11-927 (Delta) | 0.2828 | 1.4743 | 1.3307 | 2.0052 |
| TY38-873 (Omicron. BA.1) | 0.3407 | 1.6655 | 1.3846 | 1.9746 |
| TY40-385 (Omicron. BA.2) | 0.4614 | 1.0315 | 2.7310 | 3.5260 |
| **nirmatrelvir** | ($\mu\text{M}$) | ($\mu\text{M}$) | | |
| WK-521 | 1.8787 | 4.8869 | 2.5227 | 0.9858 |
| QK002 (Alpha) | 1.7795 | 6.5284 | 1.6904 | 0.7530 |
| TY7-501 (Gamma) | 1.6458 | 4.8727 | 2.0243 | 0.9244 |
| TY11-927 (Delta) | 1.7959 | 5.1161 | 2.0988 | 0.8828 |
| TY38-873 (Omicron. BA.1) | 1.8522 | 4.7245 | 2.3465 | 0.9403 |
| TY40-385 (Omicron. BA.2) | 1.9402 | 5.1784 | 2.2393 | 0.8653 |

- : The value cannot be estimated because of the low antiviral activity

### Mathematical analysis for the antibody/drug efficacy on SARS-CoV-2 variants

Based on the dose-response curves, we quantified the concentrations that achieved 50% and 90% of the maximal effect (IC_50_, IC_90_). Additionally, we estimated the Hill coefficient (m) (Koizumi et al., 2017; Shen et al., 2008), showing the steepness of the sigmoid curve, given by the equation for a fraction of infection events unaffected by drugs (f_u_),

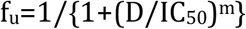

Although IC_50_ (or IC_90_) is frequently used to evaluate the “potency” of drugs, the “efficacy” of drugs is determined by both m and IC_50_: The inhibition of viral propagation (“drug efficacy”) at any given drug concentration (D) can be expressed as the instantaneously inhibitory potential (IIP) (Koizumi et al., 2017; Shen et al., 2008).

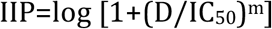

Here IIP indicates the log of viral reduction. Since antiviral drugs are usually at far higher concentrations than IC_50_ in clinical settings, the high steepness of the sigmoid curve (high m) achieves a much higher IIP than those having low steepness (low m) even if IC_50_ is the same. Hence, since sequence substitutions in the drug target [Spike, main protease (NSP5), or polymerase (NSP12)] can change IC_50_ and m, we estimated these values based on the dose-response curve for each variant through the fitting of f_u_ with nonlinear least squared regression (Table 1). To evaluate the antiviral effect in clinical drug concentrations, we calculated IIP at maximum drug concentrations [IIP(C_max_)] for each antibody/drug against each variant (Table 1 and Table S1) based on available pharmacokinetics in patients. Although casirivimab conserved IC_50_ and m (within two folds) among the Wuhan, Alpha, Gamma, and Delta strains, with profound effects to reduce viral propagation at C_max_ (3.67-7.00 log), its activity was lost to BA.1 and BA.2. While imdevimab also had a conserved antiviral effect at C_max_ (4.52-7.97 log) to the variants other than Omicron, its effect was lost on BA.1, and was retained moderately on BA.2 (2.17 log). S309’s effect at Cmax on these variants other than BA.2 was high (> 3 log), but that on BA.2 was estimated to be lower than others (1.15 log). In contrast, the IC_50_ and m for EIDD-1931 against each variant were less diverse (within two folds) to conserve strong antiviral effects at C_max_ (1.97-3.53 log). Nirmatrelvir had an even more conserved IC_50_ and m, with similar IIP at C_max_ (0.753-0.986). These analyses suggest that while the three approved-antibodies were less active to BA.1 and BA.2, EIDD-1931 and nirmatrelvir had conserved antiviral effects on variants. Our analysis suggests that the emergence of SARS-CoV-2 variants narrowed the options for efficient antibody treatments.

### Limitations of the study

This study was limited to cell culture infection assays, in which cell types and other conditions reflected drug sensitivities. However, viral targeting agents examined in this study (casirivimab, imdevimab, and S309 target Spike, EIDD-1931 target polymerase, and nirmatrelvir target main protease) were much less governed through cellular backgrounds, compared to host-targeting antivirals, and instead are more affected by viral factors such as sequence changes in the viral genome. Our assay at least compared the intrinsic antibody/drug sensitivity of SARS-CoV-2 variants side by side, helpful in discussing the impact of sequence substitutions on antibody/drug activities. Additionally, the analyses of animal and patient infections under treatment were further desired to understand drug efficacy. Yet, given the Omicron BA.2 wave’s urgency and the need for the scientific evidence to better combat this infectious disease, our data significantly present the potential diversity of drug/antibody efficacies among SARS-CoV-2 variants.

## Acknowledgments

VeroE6/TMPRSS2 cells were kindly provided by Dr. Makoto Takeda at Department of Virology III, National Institute of Infectious Diseases. This study was supported by an AMED grant JP20fk0108411 (to K.W.), Moonshot R&D Grant JPMJMS2021 (to S.I.), JPMJMS2025 (to S.I.), and JST MIRAI (to S.I. and K.W.)

## Author contributions

H.O., T.H., D.A., K.S., K.T., and K.W. performed experiments. K.S.K., J.W., S.Iwanami, and S.Iwami performed mathematical analysis. S.M., H.K., S.Y., Y.K., T.Y., N.K., S.W., H.H., H.E., T.S., K.M., S.F., and Y.T. contributed materials. S.Iwami and K.W. prepared manuscripts. K.W. supervised project.

## Methods

### Cell culture infection assay

VeroE6/TMPRSS2 cells, a VeroE6 cell clone overexpressing the transmembrane protease, serine 2 (TMPRSS2), were cultured in Dulbecco’s modified Eagle’s medium (Life Technologies) supplemented with 10% fetal bovine serum (FBS) (Sigma), 100 units/mL penicillin, 100 μg/mL streptomycin, 10 mM HEPES (pH 7.4), and 1 mg/mL G418 (Nacalai) at 37°C in 5% CO2 (Ohashi et al., 2021).

SARS-CoV-2 was investigated in a biosafety level 3 (BSL3) room. Through viral inoculation, we used WK-521 (Wuhan strain, EPI_ISL_408667), QK002 (Alpha, EPI_ISL_768526), TY7-501 (Gamma, EPI_ISL_833366), TY11-927 (Delta, EPI_ISL_2158617), TY38-873 (Omicron BA.1, EPI_ISL_7418017), and TY40-385 (Omicron BA.2, EPI_ISL_9595859) strains, isolated from COVID-19 patients and registered in GISAID. Viral infectious titers were measured by inoculating VeroE6/TMPRSS2 cells with a 10-fold serial dilution of the virus followed by measuring cytopathology to calculate TCID_50_/ml (Ohashi et al., 2021). Infection assay was conducted by inoculating VeroE6/TMPRSS2 cells with each SARS-CoV-2 strain at an MOI of 0.003 for 1 h, followed by washing out free viruses and culturing cells with fresh medium. A medium with 2% FBS without G418 was used during the infection assay. Antibodies and antiviral drugs were treated for 1 h during virus inoculation and 24 h after inoculation. At 24 h postinfection, the culture supernatant was recovered to isolate RNA using MagMax Viral/Pathogen Nucleic Acid Isolation kit (Thermo Fisher Scientific) and quantify SARS-CoV-2 RNA by real time RT-PCR with a one-step qRT-PCR kit (THUNDERBIRD Probe One-step qRT-PCR kit, TOYOBO) using 5’-ACAGGTACGTTAATAGTTAATAGCGT-3’, 5’-ATATTGCAGCAGTACGCACACA-3’, and 5’-FAM-ACACTAGCCATCCTTACTGCGCTTCG-TAMRA-3’. Simultaneously, cell viability was quantified by fixing the cells with 4% paraformaldehyde and staining with DAPI to count the number of cells using a high content imaging system ImageXpress (Molecular Devices).

