## Supplementary material for "Different efficacies of neutralizing antibodies and antiviral drugs on SARS-CoV-2 Omicron subvariants, BA.1 and BA.2": Fig. S1, Table. S1

[1]Research Center for Drug and Vaccine Development, National Institute of Infectious Diseases, Tokyo 162-8640, Japan; [2]Interdisciplinary Biology Laboratory (iBLab), Division of Biological Science, Graduate School of Science, Nagoya University, Nagoya, Japan; [3]Department of Virology II, National Institute of Infectious Diseases, Tokyo 162-8640, Japan; [4]Department of Applied Biological Science, Tokyo University of Science, Noda 278-8510, Japan; [5]Department of Virology I, National Institute of Infectious Diseases, Tokyo 162-8640, Japan; [6]Department of Veterinary Science, National Institute of Infectious Diseases, Tokyo 162-8640, Japan; [7]Center for Influenza and Respiratory Virus Research, National Institute of Infectious Diseases; Tokyo 208-0011, Japan; [8]Department of Pathology, National Institute of Infectious Diseases; Tokyo, 162-8640, Japan; [9]Institute of Mathematics for Industry, Kyushu University, Fukuoka, Japan; [10]Institute for the Advanced Study of Human Biology (ASHBi), Kyoto University, Kyoto, Japan; [11]Interdisciplinary Theoretical and Mathematical Sciences Program (iTHEMS), RIKEN, Saitama, Japan; [12]NEXT-Ganken Program, Japanese Foundation for Cancer Research (JFCR), Tokyo, Japan; [13]Science Groove Inc., Fukuoka, Japan 8100041.

[†]These authors equally contributed to this work.

*Corresponding author:
Shingo Iwami:

### Supporting Figure

**Fig. S1**

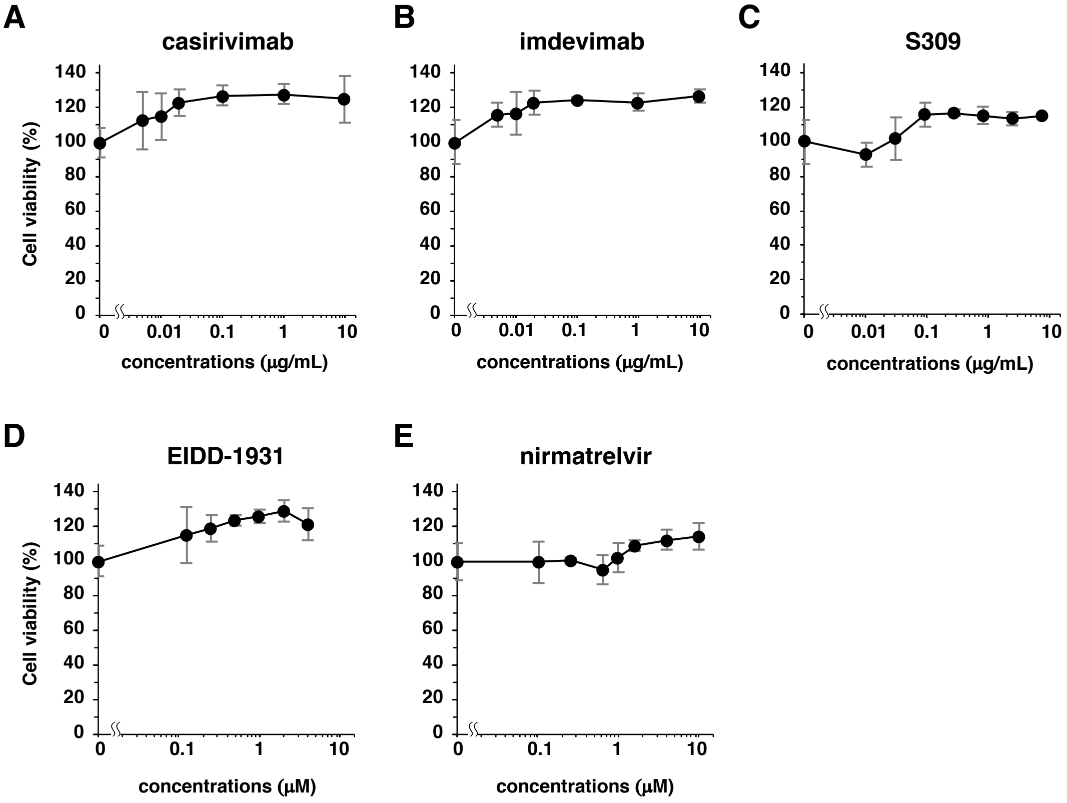


**Fig. S1. Cell viability upon treatment with neutralizing antibodies and antiviral drugs.** VeroE6/TMPRSS2 cells were inoculated with SARS-CoV-2 at an MOI of 0.003 in the presence of indicated antibodies or drugs at indicated concentrations for 1 h. The cells were then washed out and were cultured in a medium supplemented with the indicated concentrations of antibodies or drugs. After 24 h, cells were subsequently fixed with 4% paraformaldehyde and stained with DAPI, followed by counting using a high content imaging system ImageXpress Micro Confocal (Molecular Devices) to quantify cell viability. Data are presented as mean ± SD across the three replicate experiments. Relative values are shown as percentages of cell viability to the control cells that were incubated without antibodies/drugs.

### Supporting Table

**Table. S1. Drug concentrations in patients**

| Drug | $C_{max}$ (unit) | Molecular mass (unit) |
|---|---|---|
| **casirivimab** | 192 (μg/mL) | 145230 (g/mol) |
| **imdevimab** | 198 (μg/mL) | 144140 (g/mol) |
| **S309** | 117.6 (μg/mL) | 149000 (g/mol) |
| **EIDD-1931** | 2.97 (μg/mL) | 329.31 (g/mol) |
| **nirmatrelvir** | 2.21 (μg/mL) | 499.54 (g/mol) |

The pharmacokinetics of these antibodies/drugs are available as follows.

1. Food and Drug Administration (2021) Fact Sheet for Health Care Providers Emergency Use Authorization (EUA) of REGEN-COV (casirivimab and imdevimab). https://www.fda.gov/media/145611/download
2. Food and Drug Administration (2021) Fact Sheet for Health Care Providers Emergency Use Authorization (EUA) of Sotrovimab. https://www.fda.gov/media/149534/download
3. Food and Drug Administration (2022) Fact Sheet for Health Care Providers Emergency Use Authorization (EUA) of Molnupiravir. https://www.fda.gov/media/155054/download
4. Food and Drug Administration (2021) Fact Sheet for Health Care Providers Emergency Use Authorization (EUA) of Paxlovid. https://www.fda.gov/media/155050/download
